# Neuronal correlations in MT and MST impair population decoding of opposite directions of random dot motion

**DOI:** 10.1101/267732

**Authors:** Tristan A. Chaplin, Maureen A. Hagan, Benjamin J. Allitt, Leo L. Lui

**Affiliations:** Neuroscience Program, Biomedicine Discovery Institute and Department of Physiology, Monash University, Clayton, VIC 3800, Australia; ARC Centre of Excellence for Integrative Brain Function, Monash University Node, VIC 3800, Australia; Sainsbury Wellcome Centre for Neural Circuits and Behaviour, University College London, 25 Howland Street, London W1T 4JG, United Kingdom

## Abstract

The study of neuronal responses to random-dot motion patterns has provided some of the most valuable insights into how the activity of neurons is related to perception. In the opposite directions of motion paradigm, the motion signal strength is decreased by manipulating the coherence of random dot patterns to examine how well the activity of single neurons represents the direction of motion. To extend this paradigm to populations of neurons, studies have used modelling based on data from pairs of neurons, but several important questions require further investigation with larger neuronal datasets. We recorded neuronal populations in the middle temporal (MT) and medial superior temporal (MST) areas of anaesthetized marmosets with electrode arrays, while varying the coherence of random dot patterns in two opposite directions of motion (left and right). Using the spike rates of simultaneously recorded neurons, we decoded the direction of motion at each level of coherence with linear classifiers. We found that the presence of correlations had a detrimental effect to decoding performance, but that learning the correlation structure produced better decoding performance compared to decoders that ignored the correlation structure. We also found that reducing motion coherence increased neuronal correlations, but decoders did not need to be optimized for each coherence level. Finally, we showed that decoder weights depend of left-right selectivity at 100% coherence, rather than the preferred direction. These results have implications for understanding how the information encoded by populations of neurons is affected by correlations in spiking activity.

**Significance Statement:** Many studies have examined how the spiking activity of single neurons can encode stimulus features, such the direction of motion of visual stimuli. However, majority of such studies to date have only recorded from a small number of neurons at the same time, meaning that one cannot adequately account for the trial-to-trial correlations in spiking activity between neurons. Using multi-channel recordings, we were able to measure the neuronal correlations, and their effects on population coding of stimulus features. Our results have implications on the way which neural populations must be readout in order to maximize information.

## Introduction

Understanding the way in which stimulus features are represented by the activity of neurons is one of the key challenges in systems neuroscience. One of the most effective paradigms for addressing this question has been the decoding of the direction of visual motion from the activity of neurons in the middle temporal area (MT) of the primate cerebral cortex. In the classic opposite directions of motion discrimination task (Newsome et al., 1989; Britten et al., 1992, 1996), the strength of the motion signal is manipulated to reduce both the behavioural performance and the amount of information that is carried by single neurons (Newsome et al., 1989; Britten et al., 1992, 1996). While it was initially found that the activity of single neurons could account for behavioural performance (Newsome et al., 1989; Britten et al., 1992), it has become clear that the activity of a pool of neurons must be combined to form the perceptual decision (Britten et al., 1996; Shadlen et al., 1996; Law and Gold, 2008; Cohen and Newsome, 2009). Similar findings have been reported in the medial superior temporal area (MST) (Celebrini and Newsome, 1994), even though it is also associated with higher order motion processing (Saito et al., 1986; Tanaka et al., 1986; Duffy and Wurtz, 1991).

Combining the activity of a population of neurons can reduce the effects of single neuron variability (Tolhurst et al., 1983) and increase the reliability of signal. However, combining the activity of single neurons recorded on different trials, even in response to the identical stimulus, can only approximate the population responses, since it does not capture correlations in the trial to trial variability between pairs of neurons (Zohary et al., 1994; Bair et al., 2001). Correlated activity has the potential to change the amount of information contained in populations of neurons - initially, it was believed that these correlations would impair population decoding performance (Zohary et al., 1994), but it was later shown that is not necessarily the case, and that certain correlation structures can actually improve population decoding (Abbott and Dayan, 1999; Sompolinsky et al., 2001; Averbeck et al., 2006; Shamir and Sompolinsky, 2006; Ecker et al., 2011; Graf et al., 2011; Shamir, 2014; Kohn et al., 2016; Zylberberg et al., 2016). The effects of correlations on population decoding are highly dependent on the correlation structure of the population (Abbott and Dayan, 1999; Moreno-Bote et al., 2014; Kohn et al., 2016), the size of the population (Lin et al., 2015), as well as heterogeneity in firing rates and tuning (Shamir and Sompolinsky, 2006; Ecker et al., 2011; Moreno-Bote et al., 2014; Shamir, 2014; Kanitscheider et al., 2015; Kohn et al., 2016) - even untuned neurons may contribute to population coding (Goris et al., 2014; Leavitt et al., 2017; Zylberberg, 2017). Therefore, it is important to consider population activity recorded concurrently on the same set of trials to understand the information contained within a neuronal population.

Previous studies that have investigated population coding for opposite directions of motion discrimination have simulated neuronal populations by using correlation statistics that were obtained from studies of simultaneously recorded of pairs of neurons (Shadlen et al., 1996; Cohen and Newsome, 2009; Law and Gold, 2009). However, these studies did not incorporate differences correlations between different levels of motion coherences (Bair et al., 2001), and may not have incorporated the full range in of firing rates and direction tuning that are typically found in MT (Chaplin et al., 2017). More generally, it is unclear how the correlation structure of a population of (more than 2) neurons may change with motion coherence, and if this affects population coding. For example, optimal population decoding could require a different strategy depending on the strength of the motion signal, or alternatively, the optimal decoding method for high signal strength (supra-threshold) stimuli may be the same as the optimal decoding method for low signal strength (near threshold) stimuli. Furthermore, the question of whether correlations improve population decoding may depend on the level of motion coherence. Addressing these questions with data from real populations of simultaneously recorded neurons will provide valuable insights into the population activity in an opposite directions of motion task.

We recorded population activity in area MT and MST while presenting opposite directions of motion (leftwards and rightwards) and manipulated the strength of the motion signal through changes in the coherence of random dot patterns. We found that MT/MST cells were weakly correlated, and correlations increased as motion strength decreased. Correlations impaired population decoding, but decoders that accounted for the correlation structure generally outperformed decoders that ignored correlations. Despite the changes in correlation with respect to motion coherence, the same decoding strategy can be applied to all coherences with no significant loss in performance. Furthermore, decoder weights were best predicted by the individual neuron’s left-right selectivity, with some adjustments for correlations.

## Methods

### Animals and surgical preparation

Single unit and multi-unit extracellular recordings in areas MT and MST were obtained from 5 marmoset monkeys; 2 males and 3 females, between 1.5 and 3 years of age, with no history of veterinary complications. These animals were also used for unrelated anatomical tracing and visual physiology experiments. Experiments were conducted in accordance with the [National Code of Practice], and all procedures were approved by the [University Animal Ethics Committee].

The preparation for electrophysiology studies of marmosets has been described previously (Bourne and Rosa, 2003; updated as in Yu and Rosa, 2010). Anesthesia was induced with alfaxalone (Alfaxan, 8 mg/kg), allowing a tracheotomy, vein cannulation and craniotomy to be performed. After all surgical procedures were completed, the animal was administered an intravenous infusion of pancuronium bromide (0.1 mg/kg/h) combined with sufentanil (6-8 μg/kg/h, adjusted to ensure no physiological responses to noxious stimuli) and dexamethasone (0.4 mg/kg/h), and was artificially ventilated with a gaseous mixture of nitrous oxide and oxygen (7:3). The electrocardiogram and level of cortical spontaneous activity were continuously monitored. Administration of atropine (1%) and phenylephrine hydrochloride (10%) eye drops was used to produce mydriasis and cycloplegia. Appropriate focus and protection of the corneas from desiccation were achieved by means of hard contact lenses selected by retinoscopy.

### Electrophysiology, data acquisition and pre-processing

We recorded neural activity with single shaft linear arrays (NeuroNexus) consisting of 32 electrodes separated by 50 μm. MT and MST recording sites were identified during experiments using anatomical landmarks, receptive field progression and size (Rosa and Elston, 1998), and direction selectivity. The position of recording sites were confirmed post-mortem by histological examination. Penetrations were generally oriented in the dorsoventral plane, resulting an angle of approximately 30-60° relative to the surface of the cortex for MT and MST (Paxinos et al., 2012) and therefore the neuronal populations spanned both cortical columns and layers.

Electrophysiological data were recorded using a Cereplex system (Blackrock Microsystems) with a sampling rate of 30 kHz. For offline analysis of spiking activity, each channel was high-pass filtered at 750 Hz and spikes were initially identified based on threshold crossings. Units were sorted using Offline Sorter (Plexon Inc.). Units were classified as single-units if they showed good separation on the (2 component) principal component analysis plot, and were confirmed by inspection of the interspike interval histogram and consistency of waveform over time. Any remaining threshold crossings were classified as multi-unit activity. We used only one unit from each electrode contact – either the best isolated single unit, or the multiunit if there were no single units (which excluded 23 responsive units). We also excluded 9 units from adjacent channels since it was apparent they were duplicated across two channels, based on their sharp cross correlogram peak and high signal and noise correlations (Bair et al., 2001). To account for the possibility that we may have still double counted spikes across adjacent channels, we also removed coincident spikes on adjacent channels using the following procedure. We compared the spike train of each responsive unit to the unit at the immediately adjacent electrode contact (50 μm away), and deleted any spikes in the second unit’s spike train that occurred within plus or minus 1ms of spikes in the first unit, thus eliminating any double counted spikes (but also removing some that occurred at the same time by chance). This eliminated a further 28 out of 221 units from the dataset, because they were now considered unresponsive to the best stimulus (compared to the spontaneous rate, Wilcoxon rank sum test p < 0.01, see the Data Analysis section below).

### Visual stimuli

Visual stimuli were presented on a VIEWPixx3D monitor (1920 × 1080 pixels; 520 × 295 mm; 120 Hz refresh rate, VPixx Technologies) positioned 0.35 to 0.45 m from the animal on an angle to accommodate the size and eccentricity of the receptive field(s), typically subtending 70° in azimuth, and 40° in elevation. All stimuli were generated with MATLAB using Psychtoolbox-3 (Brainard, 1997).

The main visual stimulus consisted of random dots presented full screen. White dots (106 cd/m^2^) of 0.2° in diameter were displayed on a black (0.25 cd/m^2^) background (full contrast). The density was such that there were on average 0.5 dots per °^2^; this was chosen because these parameters have been shown to elicit good responses from marmoset MT when displayed on LCD monitors (Solomon et al., 2011; Zavitz et al., 2016). Dot coherence was controlled using the white noise method (i.e. Britten et al., 1992, 1996; see Pilly and Seitz 2009) by randomly choosing a subset of “noise” dots on each frame, which were displaced to random positions within the stimulus aperture. The remaining “signal” dots were moved in the same direction with a fixed displacement. For each stimulus presentation, a new set of signal and noise dots were randomly generated, thereby producing a different stimulus for each trial.

#### Determination of receptive fields and basic direction tuning

Visual receptive fields were quantitatively mapped using a grid of either static flashed squares or small apertures of briefly presented moving dots. Subsequent visual stimuli were presented full screen, so as to cover as many neurons’ receptive fields as possible. We also conducted visual direction tuning tests (12 directions, 100% coherence), sometimes using just a single standard speed of 20°/s, other times a range of speeds to determine the optimal speed. Units were deemed to be direction selective if their circular variance (Ringach et al., 2002) was less than or equal to 0.9 (cutoff determined by visual inspection of the direction tuning curves and the distribution of the circular variances). The preferred direction was calculated with a vector sum (Ringach et al., 2002), and in analyses that used preferred direction, we only used units with a circular variance less than or equal to 0.9 (i.e. were direction selective). The strength of the direction selectivity was calculated with a direction index DI:

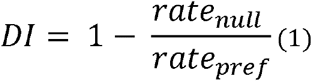

Where rate_pref_ and rate_null_ is the firing rate in response to preferred and null direction of motion respectively, at 100% motion coherence. DI values lie between 0 and 1, with 1 indicating a strongly direction selective neuron.

#### Main stimulus protocol

As our investigations involved decoding for opposite directions of motion, we presented visual stimuli moving either leftwards or rightwards at 60°/s at different levels of motion coherence: 100, 82, 64, 46, 28, 10 and 0%. All stimuli were presented for 600 ms with 120 repeats per condition. The number of trials per direction and coherence meant that this test protocol took up considerable time, therefore we collected data from only one axis of motion to maximize recording stability. There was no specific reason why the left-right axis was chosen, apart from the fact that these data were obtained as part of a study to investigate the effects of auditory motion in MT and MST. Because we did find any effect of auditory stimuli in the responses of single neurons or populations of neurons, we have grouped the two conditions (visual and audio-visual) into one dataset for these analyses. As we used the horizontal axis of motion regardless of the direction preferences of the recorded units, our results will be generalizable to the opposite directions paradigm across all directions.

### Data Analysis

#### Time windows and inclusion criteria

Firing rates were calculated using a time window from the stimulus onset to offset (600 ms). Units were deemed responsive if the firing rate in response to the best 100% coherence stimulus (leftwards or rightwards) was significantly different to the spontaneous rate (Wilcoxon rank sum test, p < 0.01), and if this rate was at least 2 spikes/s above the spontaneous rate. Units were considered left-right selective if the firing rate to the best direction of motion (leftwards or rightwards) at 100% coherence was significantly greater than that of the other direction (Wilcoxon rank sum test, p < 0.05). Populations were included for analysis if they contained at least 3 left-right selective units that were separated by at least 100 μm. When comparing the results across the main stimulus protocol and the direction tuning protocol (Figures 2C and 6E), we excluded 10 units due to significant changes in left-right selectivity, possibly due to small movements in the electrode, and 19 units that were not responsive at the time of the direction tuning test. For these analyses that used preferred direction, we only used units that were direction selective (CV <= 0.9), because the preferred direction is meaningless for non-direction selective units.

#### Left-right selectivity

We characterized the left-right selectivity of each unit with a left-right index:

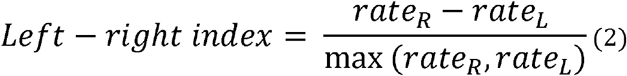

Where rate_R_ and rate_L_ is the firing rate in response to rightwards and leftwards motion respectively, at 100% motion coherence. Left-right indices lie between −1 and +1, with −1 indicating a strongly leftwards preferring neuron, +1 indicating a strongly rightwards preferring neuron, and 0 indicating a neuron that is not selective for leftwards or rightwards motion. The left-right congruency of a pair of units was defined as the product of the two left-right indices, resulting in a range of values from −1 to +1, where −1 is a pair of strongly left-right selective units that have opposite left-right preferences, and +1 is a pair of strongly left-right selective units that have the same left-right preference.

#### Decoding

We used Linear Discriminant Analysis (Pesaran et al., 2002; Averbeck et al., 2003; Law and Gold, 2009; Adibi et al., 2014) to decode the direction of motion (leftwards or rightwards) at each coherence. Firing rates were first z-scored at each coherence before being used for decoding, using the mean and standard deviation of the combined firing rates to leftwards and rightwards motion. We used random subsampling cross validation by training on a randomly selected subset of 80% (96/120) of trials and testing on the remainder, and repeating this process 1000 times. For each iteration, we trained and tested the decoder at each level of motion coherence. Therefore, we obtained an estimate of the decoding accuracy (the mean percent correct across iterations) and the variability (95% interval across iterations). We also applied the same method to decode the direction of motion from each individual unit in order to compare them to the population performance. Since the range of weights varied by population, weights were normalized by dividing by the maximum absolute weight in the population when comparing across populations.

#### Neurometric thresholds

To determine neurometric thresholds of single neurons and populations (Newsome et al., 1989; Britten et al., 1992; Berens et al., 2011), the percent correct values of each cross-validation iteration were fitted using least squares regression with two variants of the Weibull function, resulting in a neurometric curve that described the decoding performance with respect to coherence:

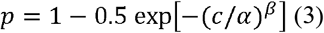

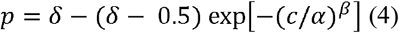

where *p* was the probability of correctly discriminating the direction of motion at coherence *c, α* was the coherence of threshold performance (82%, convention established by Britten et al., 1992), β controlled the slope and δ was the asymptotic level of performance (less than 1). As Equation 4 has an extra free parameter, we used an F-test to decide whether to reject the use of Equation 4 over Equation 3. The α was limited to between 0 and 3, β was limited to lie between 0 and 10, and δ was limited to lie between 0 and 1. In some analyses, we used the “near-threshold” level of coherence, which was the coherence that was closest to the exact threshold as determined by the curve fitting procedure. We only analyzed populations in which the upper bound of the threshold’s 95% interval was less than 100% coherence, in order to ensure that threshold estimates were well constrained. Therefore, all populations had a population threshold less than 100% coherence.

#### Decoding with and without correlations

To test the effects of correlations on population decoding, we trained two types of decoders; the standard decoder, which was trained on the standard dataset (i.e. contains correlations), and a “correlation blind” decoder, which was trained on trial shuffled datasets, a process which removed all correlations. To test the effect of removing correlations, we compared the performance of the blind decoder on trial shuffled dataset to the standard decoder on the standard dataset. To test the effect of ignoring correlation structure, we compared the performance of the blind decoder to the standard decoder on the standard dataset, i.e. a data set that contained real correlations.

#### Spike count correlation

For each pair of units in each population, we calculated the spike count correlation (r_SC_) as the Pearson’s correlation coefficient of the trial by trial spike counts. We calculated rSC for each pair of units by z-scoring the firing rates at each direction and coherence, so the firing rates could be combined across stimulus conditions to calculate an overall rSC for the pair. We also calculated rSC for each level of motion coherence and direction separately. The rSC values were calculated for each iteration of the random subsampling cross validation procedure used for decoding and averaged across iterations.

#### Statistics

Measures of correlations were Spearman’s rho (p), except for spike count correlations (see above). Tests between two groups were made with Wilcoxon’s Rank Sign test (paired) or Wilcoxon’s Rank Sum test (unpaired), and the α criterion was 0.05 unless otherwise specified. Running means were calculated using 1000 sliding windows. The window size was 0.15 and 10° for left-right congruency and difference in preferred direction respectively. For the electrode separation, we only plotted the running mean for separations that had least 10 pairs of units, and for left-right congruency, we only plotted for the middle 95% percentile to ensure a reliable estimate of the mean. To test for differences in spike count correlations at different levels of motion coherences, we used a 2-way ANOVA with the Tukey-Cramer method for post hoc multiple comparisons, and we used an ANCOVA to account for differences in spike count correlations that might rise from differences in firing rate. We deemed differences in decoding thresholds to be statistically significant if the 95% interval of differences across cross-validation iterations did not overlap with zero.

### Histology

At the end of the recordings, the animals were given an intravenous overdose of sodium pentobarbitone and, following cardiac arrest, were perfused with 0.9% saline, followed by 4% paraformaldehyde in 0.1 M phosphate buffer pH, 7.4. The brain was post-fixed for approximately 24 hours in the same solution, and then cryoprotected with fixative solutions containing 10%, 20%, and 30% sucrose. The brains were then frozen and sectioned into 40 μm coronal slices. Alternate series were stained for Nissl substance and myelin (Gallyas, 1979). The location of recording sites was reconstructed by identifying electrode tracks and depth readings recorded during the experiment. Additionally, each electrode array was coated in DiI, allowing visualization under fluorescence microscopy prior to staining of the sections. In coronal sections, MT is clearly identifiably by heavy myelination in the granular and infragranular layers (Rosa and Elston, 1998), whereas MST is more lightly myelinated and lacks clear separation between layers (Palmer and Rosa, 2006). The majority of neurons reported here were histologically confirmed to be in MT or MST, but for some penetrations in which the histology was unclear (2 penetrations), neurons were included on the basis of their receptive field size and progression, and their direction tuning.

## Results

### Sample size

We made 27 electrode array penetrations in areas MT and MST, but restricted our analysis to 17 populations (see Methods for inclusion criteria, MT: n = 13; MST: n = 4) that were suitable for population decoding. The number of units per population varied from 4 to 27 (median = 11, total = 193 across all populations), comprising of both single (12%) and multi-units, but no distinctions were made between unit type for population decoding. We first performed direction tuning tests using 100% coherence motion in 12 equally spaced directions (Figure 1A-C, left column), and found that most units were direction selective (MT 80%, MST 70%, circular variance <= 0.9), in agreement with previous reports (Zeki, 1974; Maunsell and Van Essen, 1983; Albright, 1984; Celebrini and Newsome, 1994; Born and Bradley, 2005; Lui and Rosa, 2015). Cells in both areas have been shown to respond in a direction selective way to random dot stimuli (Britten et al., 1992; Celebrini and Newsome, 1994), and since decoding performance and direction selectivity in MT was similar to that of MST, we did not distinguish between MT and MST in this paper. Since the electrode penetrations were not perpendicular to the surface of the cortex, they spanned a number of direction columns and typically showed a range of different preferred directions in each population. Overall, the full set of units across all populations covered the full range of preferred directions and varying degrees of direction selectivity (Figure 1D, left panel).

**Figure 1:**
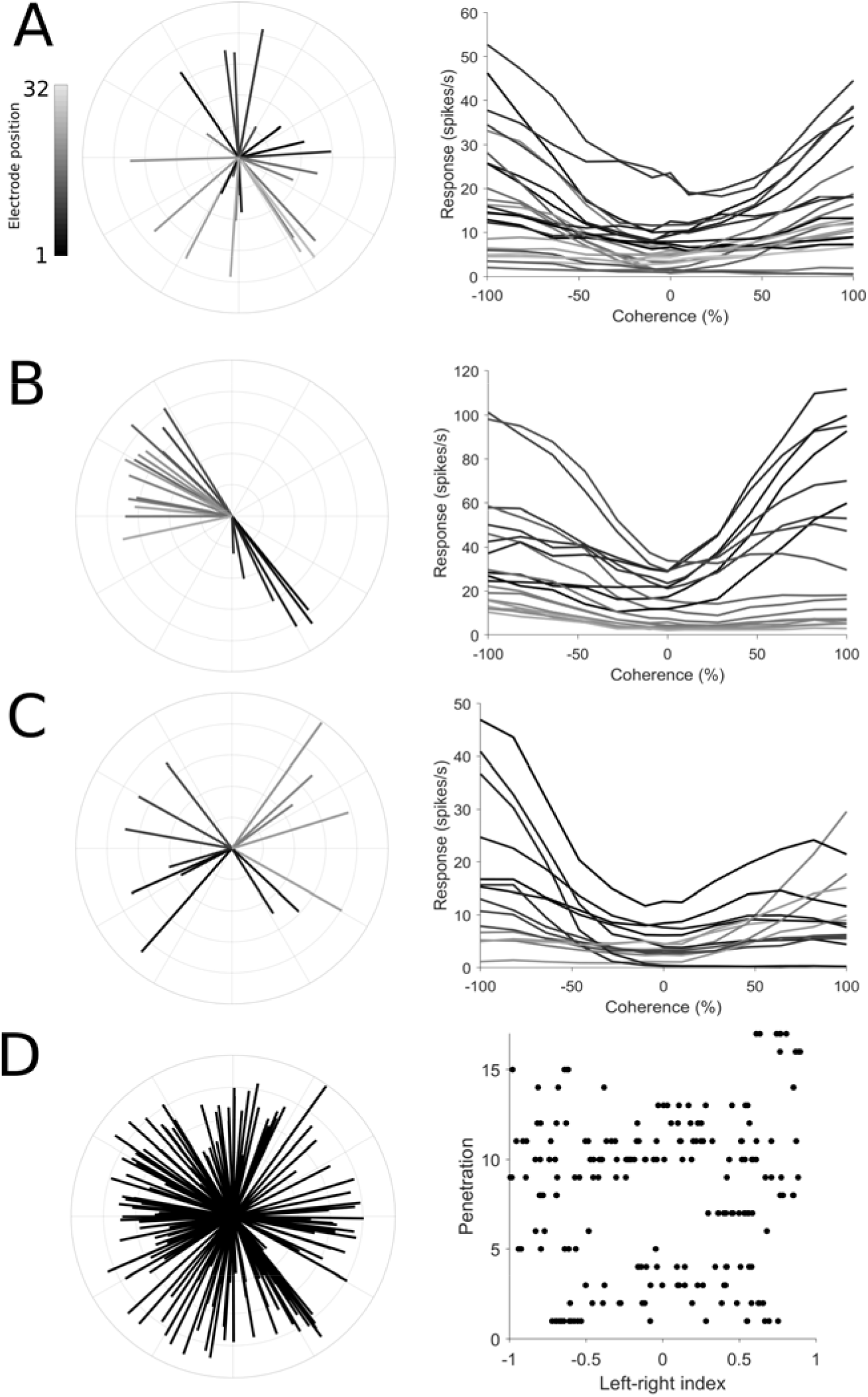
Direction preference and left-right selectivity of the recorded neuronal populations. Panels A-C show the direction tuning and the responses to left-right motion at difference coherences for 3 example populations (A and B are in area MT, C is in area MST). In the left column’s polar plots, the direction of each line represents the preferred direction of a unit, the length represents the strength of the direction selectivity (direction index, Equation 1), and the shading shows the position of the unit on the electrode shank. In the right column’s Cartesian plots, the mean spiking rate to motion on the left-right axis of motion are plotted at different levels of motion coherence. Each line shows the coherence response function for a unit from the polar plot shown in the left column, using the same shading convention. Leftwards motion is represented by negative coherence values, rightwards motion by positive coherence values. D: Distribution of preferred directions and left-right selectivity across all populations in this study. The left panel’s polar plot shows the combined set of preferred directions (same convention as A-C) for all recorded units. In the right panel, each row of dots represents a population, and the position of each dot along the x-axis represents the left-right index (Equation 2). Dots positioned on the left (negative) side of the axis prefer leftwards motion, dots positioned on the right (positive) side of the axis prefer rightwards motion.

For the main stimulus protocol in this study, we presented motion in the left-right axis at various levels of motion coherence. We observed a wide range of firing rates in responses to changes in motion coherence along the left right-axis of motion. Some units showed monotonic increases in firing from left to right or vice versa (i.e. were strongly left-right selective), other units showed “u-shaped” type responses, where firing rates increased with coherence in both directions (Figure 1A-C, right column). More often than not, these cells had preferred directions which were far away from the horizontal axis. Overall we observed a range of left-right selectiveness (Figure 1D, right panel), with most populations showing a mixture of left and right preferring units.

### The spiking activity of MT and MST neurons is weakly correlated

We first characterized the spike count correlations (r_SC_) of pairs of units in our populations. We measured rSC of all pairs of units (n = 1466) by z-scoring the firing rates for each coherence and direction and collapsing across all conditions. Confirming previous reports, we found that the activity of many pairs MT and MST neurons were weakly positively correlated on a trial to trial basis (Zohary et al., 1994; Bair et al., 2001; Cohen and Maunsell, 2009; Solomon et al., 2015; Ruff and Cohen, 2016; Zavitz et al., 2016), with a mean r_SC_ of 0.23 (Figure 2A, left panel). Because it had been previously reported that rSC depends the distance between neurons and the differences in their preferred directions (Cohen and Newsome, 2008; Solomon et al., 2015), we also investigated these factors in our dataset. We found that there was a significant negative correlation between rSC electrode separation (Figure 2A, right panel, Spearman’s ρ = 0.278, p < 0.001), left-right congruency (the product the left-right indices, Equation 2, Figure 2B, Spearman’s ρ = 0.101, p < 0.001) and the difference in preferred direction of the pair (Figure 2C, Spearman’s ρ = 0.114, p = 0.001), demonstrating that pairs of units that are closer to each other, or have similar direction tuning, tend to have higher correlations.

**Figure 2:**
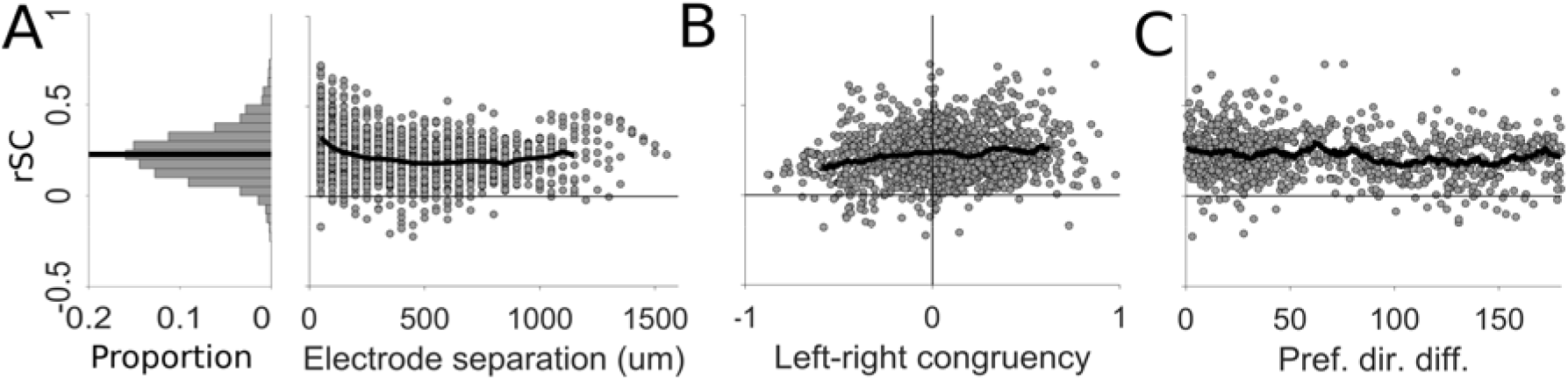
Factors that affect spike count correlations (r_SC_). A: The histogram on the left shows the distribution of r_SC_ in the dataset, and black line indicates the mean. The scatter plot on the right shows the r_SC_ of each pair plotted against electrode separation, and the black line shows the mean r_SC_ at each separation (only plotted for electrode separations that had at least 10 pairs of units). B: r_SC_ plotted against left-right congruency (the product the left-right indices, Equation 2), the black line shows the running mean (bin size = 0.15, only plotted for the middle 95 percentile range, in order to obtain reliable estimate of the mean). C: r_SC_ plotted against the difference in preferred direction of pairs of direction selective units, black line indicates the running mean (bin size = 10°).

### Low coherence stimuli produce stronger spike count correlations

Because the primary stimulus manipulation in this study was motion coherence, we next tested if r_SC_ varied with this parameter. We found that there was a statistically significant modulation of rSC by coherence (repeated measures ANOVA F12 = 105, p < 0.001) across the 13 levels of coherence (6 levels leftwards, 6 rightwards, and the zero coherence condition). Because we were primarily interested in the modulation of rSC by coherence rather than direction, we grouped the responses to leftwards and rightwards motion, and plotted the mean r_SC_ at each coherence (Figure 3A, solid line), and observed a clear effect: rSC was lower for 100% coherence compared to all other coherence levels. As the non-zero coherence levels now had two rSC values per pair (leftwards and rightwards) and the zero coherence only had one rSC value per pair, we first analyzed non-zero coherences only in order to perform repeated measures ANOVAs with respect to coherence. Confirming the previous findings, we found that r_SC_ was significantly modulated by non-zero coherences (repeated measures ANOVA F_5_ = 300, p < 0.001), and that lower coherences had higher r_SC_ measurements (post hoc tests of 100% vs 10, 28, 46, 64 and 86% coherence all p < 0.001), and that there was a significant negative correlation between coherence and r_SC_ (Spearman’s ρ = −0.15, p < 0.001). In summary, these results suggest that reducing motion coherence increases spike count correlations.

**Figure 3:**
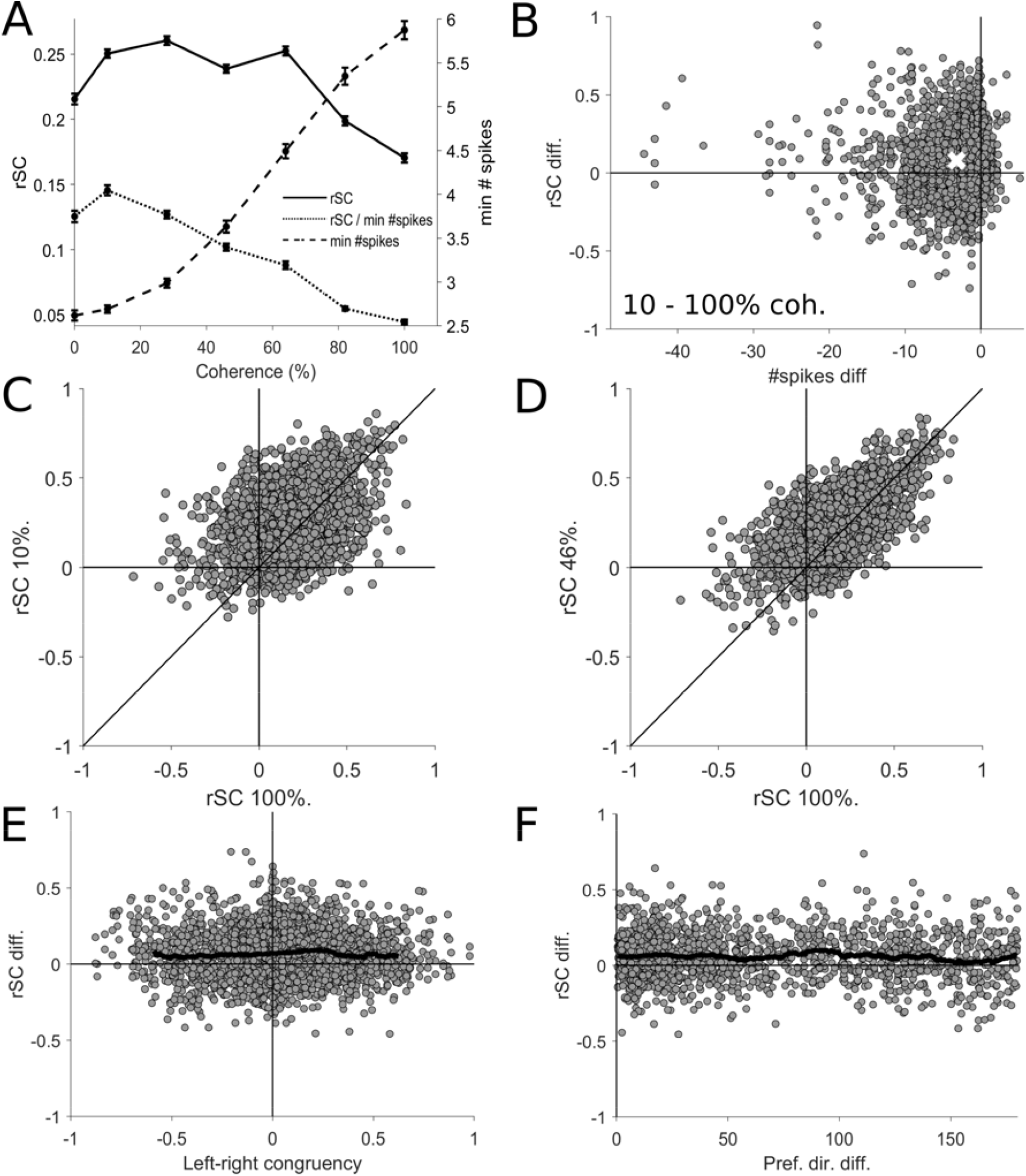
Spike count correlation (r_SC_) at different levels of motion coherence. A: The solid line shows the mean r_SC_ plotted for each level of coherence, demonstrating that low coherences produces higher r_SC_ values (left axis) than 100% coherence. The dashed line shows the mean minimum spike count (right axis) at each coherence, demonstrating that spiking activity increases with coherence, which could cause a trivial increase in r_SC_, not the observed decrease (solid line). The dotted line shows the mean r_SC_ normalized by dividing by the minimum spike count of the neuronal pair plotted against each level of coherence, confirming that the decrease in r_SC_ with coherence is not trivially due to decreases in spiking activity. Error bars show the standard error of the mean. B: The difference in r_SC_ and minimum spike count for each pair of units at 10 and 100%, the white cross indicates the means. Note the r_SC_ values were higher at 10% coherence but the spike counts were lower. C and D: r_SC_ values at 100% coherence are shown with respect to the r_SC_ values at lower coherences. E: The difference in r_SC_ for 46 and 100% coherence plotted against left-right congruency, showing no relationship. The black line shows the running mean (bin size = 0.15, only plotted for the middle 95 percentile range). C: The difference in r_SC_ for 46 and 100% coherence plotted against the difference in preferred direction of pairs of direction selective units, showing no relationship. The black line indicates the running mean (bin size = 10°).

However, such a change in r_SC_ could be trivially explained by a change in the number of spikes elicited by different motion coherences (de la Rocha et al., 2007; Cohen and Kohn, 2011), and changes in coherence clearly modulate spiking activity (Figure 1A-C right panels, Figure 3A dashed line, Britten et al., 1993; Chaplin et al., 2017). To investigate whether the observed increase in rSC with at lower coherences was simply due to changes in the spike counts, we first used an ANCOVA controlling for the effect for the minimum spike count of the pair (Cohen and Kohn, 2011) and still found a significant effect of coherence on rSC (ANCOVA F5 = 14.2, p < 0.001, post hoc tests for 100% vs 10, 28, 46, and 64% coherence p < 0.001, 100% vs 82% p = 0.91). Additionally, we normalized r_SC_ by dividing by the minimum spike count of each pair and investigated this measure against coherence, and found that the strength of normalized correlations still decreased with coherence (Figure 3A dotted line, repeated measures ANOVA F5 = 358, p < 0.001, post hoc tests for 100% vs 10, 28, 46, 64 and 86% coherence all p < 0.001). Furthermore, we found that the normalized correlations were significantly negatively correlated with coherence (Spearman’s ρ = −0.37, p < 0.001). To visually confirm that the differences in rSC between coherences and confirm they were not caused by changes in spike count, we examined the difference in rSC against the difference in spike counts for all pairs of coherence (e.g. 10 vs 100% coherence shown in Figure 3B). When comparing the difference in rSC between coherences, we found that the increases in rSC were actually accompanied by a *decrease* in spike count (e.g. Figure 3B has most points on the left side of the y-axis), thereby demonstrating that the increase in r_SC_ was not simply caused by an increase in the number of spikes. To test if the finding that 0% coherence produced lower values of rSC compared to 10% coherence because differences in spike counts (Figure 3A), we used ANCOVAs to control for minimum spike count and performed post hoc tests and found no significant effect (p = 0.935 0% vs leftwards 10% coherence motion; p = 0.875 0% vs rightwards 10% coherence motion). Therefore, these analyses demonstrate that reducing motion coherence increases spike count correlations independent of changes in spiking activity.

Finally, we examined whether correlations between pairs of neurons remained consistent between coherences, or if r_SC_ were highly variable between pairs, resulting in a different correlation structure from one coherence to the next. We examined the r_SC_ at each coherence against the r_SC_ at 100% coherence (e.g. Figure 3C-D) and found strong and significant correlations (100 vs 10, 28, 46, 64 and 86% coherence Spearman’s ρ = 0.381, 0.474, 0.585, 0.601 and 0.733 respectively, all p < 0.001). This suggests that at least some parts of the correlation structure may be preserved across coherences, even though the magnitude of the correlations changed (Figure 3A). We also examined if the change in r_SC_ with coherence was affected by the direction preference of the pairs of neurons, but did not find any systematic significant relationships with left-right congruency (e.g. Figure 3E, 100 vs 46% coherence Spearman’s ρ = 0.017, p = 0.372) or difference in preferred direction (e.g. Figure 3F, 100 vs 46% coherence Spearman’s ρ = −0.033, p = 0.207)

### Population decoding always outperforms the best unit

Having characterized the correlations in our populations of MT and MST units, we next decoded the direction of motion (leftwards or rightwards) by training and testing linear decoders at each level of motion coherence using the spike rates of each unit. The method that we used here ensured that the decoder took into account the correlations during training, and were tested on real trials where cell-to-cell covariations remained intact. We calculated a neurometric threshold for each population, defined as the level of coherence in which the decoder achieves 82% accuracy. First, we needed to confirm that decoding the direction of motion from a population was in fact incorporating the information from multiple units, and not relying solely on the single most informative unit. In order to assess the improvement in decoding using a population of neurons over an individual neuron, we also decoded the direction of motion (and calculated thresholds) for each unit individually. Figure 4A shows the decoding performance of a representative population with the population decoding performance shown in black and the best individual unit decoding performance in shown in grey. In this population, the population threshold is lower than the best unit’s threshold. In fact, across the full set of populations, the population threshold was always lower than the threshold of the best unit (Figure 4B, all points lying bellow the line of unity, median difference = −17%, p < 0.001, Wilcoxon Rank Sign test). This demonstrates that the population decoding method we used is incorporating information from multiple neurons, and that the populations as a whole contained more information about the direction of motion than any individual unit. Furthermore, the vast majority of decoding weights were non-zero (Figure 6D-E), demonstrating that most units make at least some contribution to the decoding outcome.

**Figure 4:**
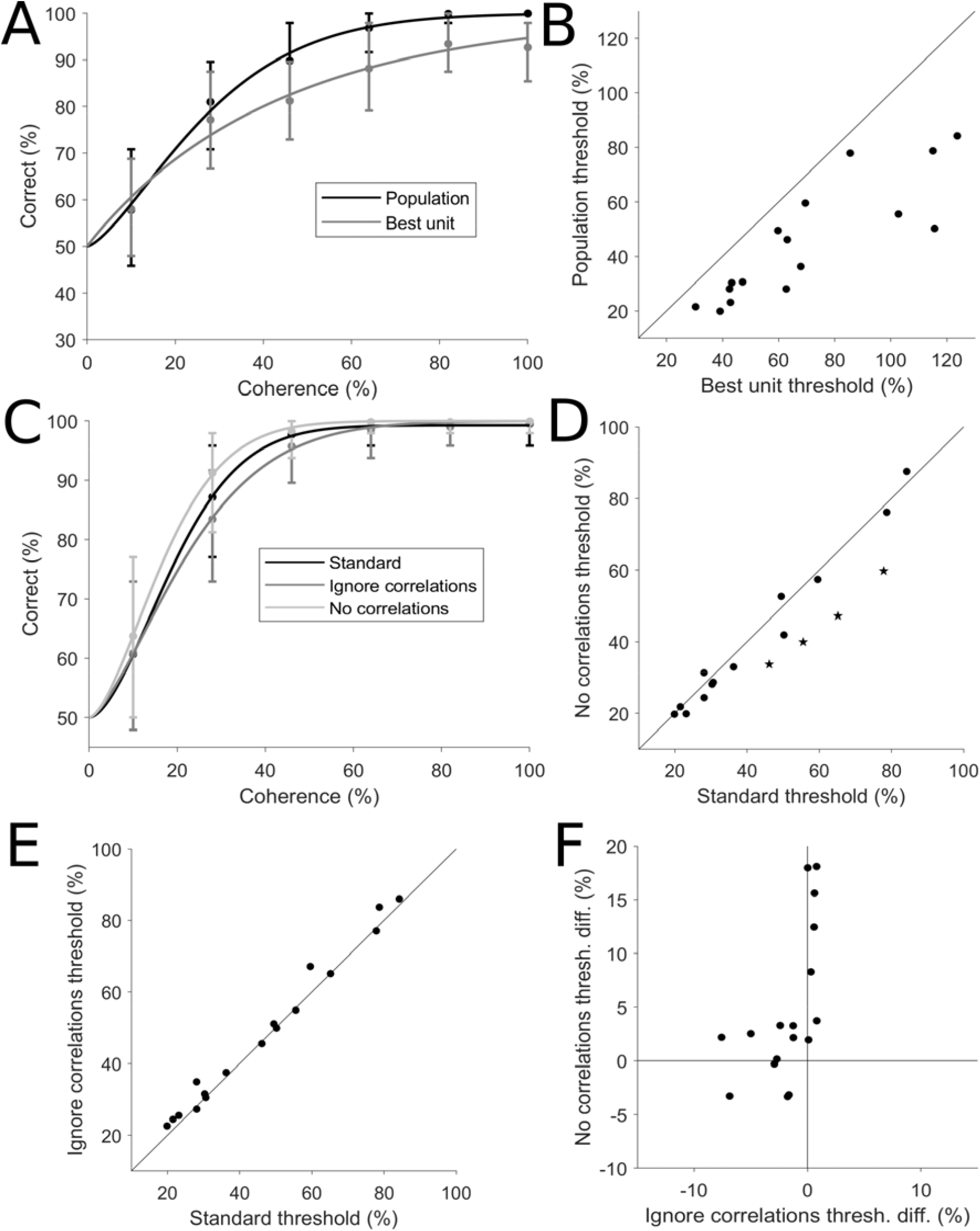
Population decoding and correlations. A: An example population showing the decoding performance of the population (black) and the best individual unit (grey) plotted against coherence. The two data sets were both fitted with a Weibull curve to determine the threshold, defined as the level of coherence that achieves 82% correct. B: Population thresholds plotted against the threshold of the best unit for all populations. All populations had thresholds lower than the threshold of the best individual unit. C: Another example population, showing the performance of the standard decoding procedure (black), performance when correlations were removed (light grey) and performance when correlations were ignored (dark grey). D: Effects of correlations. The thresholds from the standard decoding procedure are plotted against the thresholds obtained when correlations were removed, showing a statistically significant decrease in the median threshold (p = 0.025, Wilcoxon Rank Sign test). Star symbols represent populations in which the difference is statistically significant (p < 0.05; bootstrap). E: Effects of ignoring correlations. The thresholds from the standard decoding procedure are plotted against the thresholds obtained when correlations were ignored, showing a small but statistically significant increase in the median threshold (p = 0.007, Wilcoxon Rank Sign test). F: The change in thresholds when correlations were ignored was significantly positively correlated with change in thresholds when correlations were removed (Spearman’s ρ = 0.647, p = 0.005). The change in threshold was calculated as the standard threshold minus the remove or ignore correlations threshold. Therefore, for the x-axis, negative values represent populations which had lower (better performance) thresholds in the standard decoding compared to decoding when correlations where ignored. These were the units that were above the line of unity in E, and were the majority of populations. For the y-axis, positive values indicate populations that had higher thresholds (worse performance) for the standard decoder in comparison to decoding when correlations where removed. These were the units below the line of unit in D and were the majority of the populations.

Finally, we examined the factors that influence the population threshold. The threshold of the best unit was the strongest predictor of the population threshold (Spearman’s ρ = 0.9, p < 0.001). There was also a significant relationship between population threshold and the summed absolute left-right of the population, even when controlling for the threshold of the best unit (Spearman’s ρ = −0.523, p = 0.0352), demonstrating that populations with more strongly direction selective neurons have lower population thresholds. There was no significant relationship between the population threshold and the population size (Spearman’s ρ = −0.248, p = 0.337) or the mean absolute left-right selectivity (Spearman’s ρ = −0.37, p = 0.144).

### Correlations impair decoding performance

Having previously established the basic characteristics of trial-to-trial correlations, we next examined how these correlations affected population decoding. This was done by comparing the performance of the standard decoder (trained and tested with correlations present) to the trial shuffled decoders (trained and tested with no correlations present). Figure 4C shows the performance of a single example population with and without correlations (black vs light grey lines respectively), in which decoding performance was worse in the presence of correlations. In general, we found the presence of correlations impaired decoding performance across the full dataset (Figure 4D, median threshold difference = −2.7%, p=0.025, Wilcoxon Rank Sign test), and four populations showed statistically significant higher thresholds in the presence of correlations (bootstrap, p < 0.05). Therefore neuronal correlations resulted in a decrease in population decoding performance for opposite directions of motion.

We next investigated if knowing the correlation structure provided any advantage, or if similar performance could be achieved by ignoring correlations. To test this, we the compared the performance of decoding the real data set (correlations present) with the standard decoders (trained with correlations present) and trial shuffled decoders (trained with no correlations present). The example population in Figure 4C show an improvement in decoding performance when correlations were considered (black vs dark grey lines). While no individual population showed a significant decrease to threshold when taking correlations into account, there was a significant decrease in median threshold across populations when the decoder was trained with the correlation structure intact (Figure 4E, median difference = 1.6%, p = 0.007, Wilcoxon Rank Sign test). Therefore, decoders which learnt the correlation structures usually performed better than one that ignored correlations.

Next, we investigated if there was any relationship in the effect size (and sign) of removing and ignoring correlations when decoding our dataset of neuronal populations. One may expect that the two are tightly coupled, i.e. for populations that were most affected by the presence of correlations, it may be most advantageous for the decoder to learn these correlations. Such a relationship would result in a negative correlation between the effect of removing and ignoring correlations, with data points occupying the first (top left) and the third (bottom right) quadrants of Figure 4F. However this was not the case - there was in fact a significant positive correlation (Figure 4F, Spearman’s ρ = 0.647, p = 0.005). Interestingly, for the populations that were most affected by the presence correlations, ignoring correlations had little or no effect on the accuracy of decoding. Correspondingly, for the populations in which learning correlations most improved decoding performance, removing correlations did not affect decoding performance. These results were exemplified by the fact that the majority of data points in Figure 4F were situated close to both axes. In summary, our data show that the effects of removing and ignoring correlations in population decoding were related - populations that were affected by one of these factors were less likely to be affected by the other.

### Decoders trained at 100% coherence generalize to other coherences

We had previously found that spike count correlations decrease with motion coherence (Figure 3A), yet the correlation structure appeared to be preserved across coherences - e.g. pairs of units that are strongly correlated at 100% coherence are also strongly correlated at 10% coherence (Figure 3C-D). It was therefore unclear if the changes in the strength of the correlations with coherence would affect the optimal linear decoding strategy – i.e. is it necessary for the decoder to be optimized for each coherence level by training on responses from each individual level of coherence? Or alternatively, can a decoder trained on 100% coherence be successfully applied to lower coherences? If the former case is true, then downstream areas that decode MT/MST neurons would first need to be aware of the level of motion coherence (i.e. in order to use a decoding strategy that is optimized for the particular coherence), but if is the latter, then the downstream areas could always use the same readout method.

To investigate this, we used the decoders trained at 100% coherence to decode the direction of motion at every other level of coherence, and thereby obtained a new set of population thresholds, and we then compared these new thresholds to the thresholds obtained by training at each coherence level. We found there was no significant difference in population threshold between these two training methods (Figure 5A, median threshold difference = −0.53%, p = 0.717, Wilcoxon Rank Sign test), suggesting that the readout method for 100% coherence can generalize to all other levels of motion coherence. Further, we tested if decoders trained at 100% coherence do indeed perform as well as coherence specific decoders at all levels of coherence (not just near threshold), we plotted the difference in accuracy for the two decoder types at each level of coherence (Figure 5B). Firstly, we found that the difference in performance between the two decoder types was not significantly different to zero for all coherence levels (uncorrected t-tests, p < 0.05). We also found that there was no difference between coherence levels (1-way repeated measures ANOVA F4 = 1.61, p = 0.18). These results confirm that decoders trained at 100% coherence perform as well as coherence specific decoders at all levels of coherence. To directly compare these decoding methods, we compared the normalized weights of the decoders trained at 100% coherence with the decoders trained at the near threshold level of coherence (the closest level of coherence to the populations’ threshold, therefore varying by population) by plotting one against another and found they were highly correlated (Figure 5C, Spearman’s ρ = 0.8, p< 0.001), and that most weights are not significantly different under the two conditions (151 out 193 units show no significant difference with bootstrapping).

**Figure 5:**
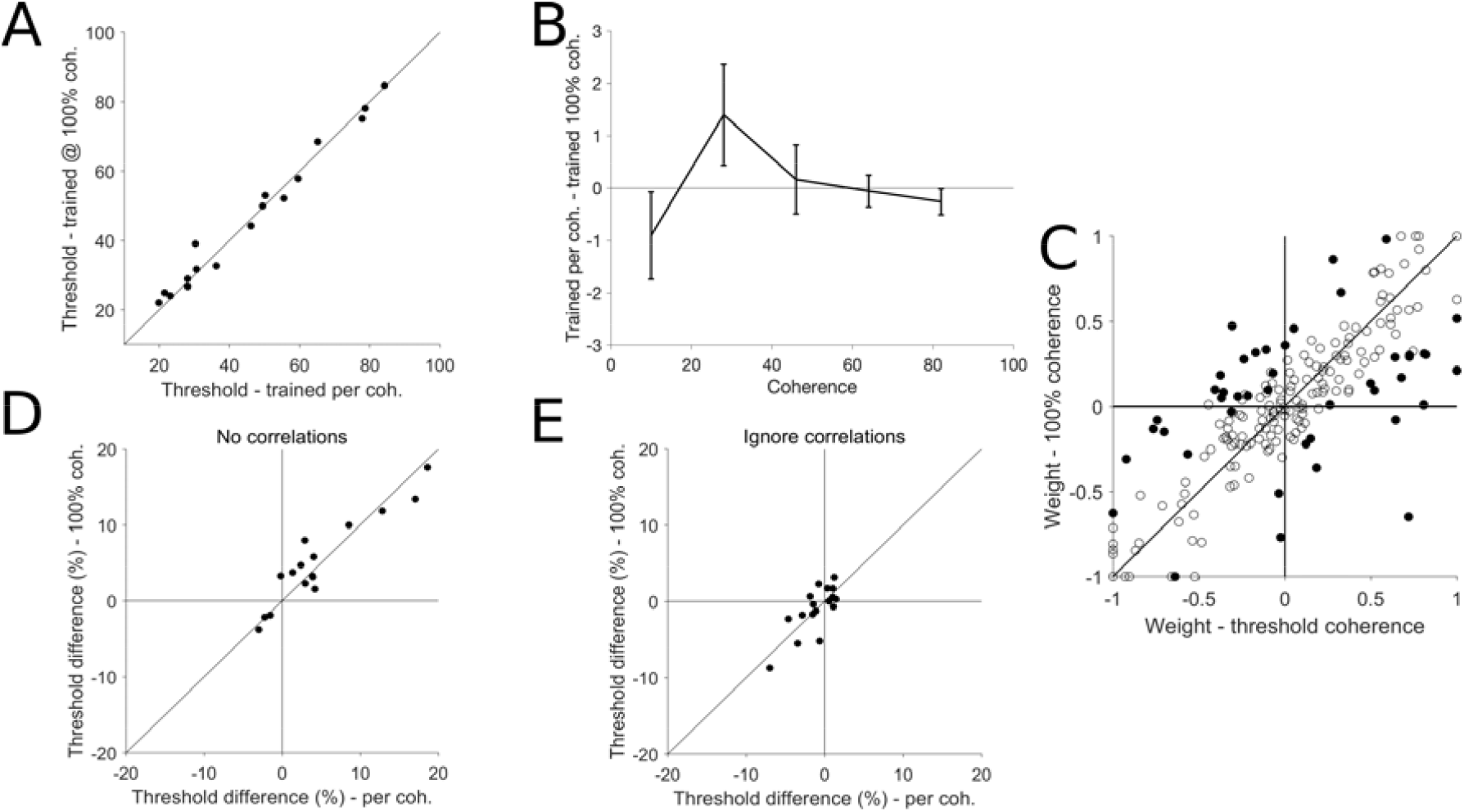
Generalizability of decoders across motion coherence. A: Population thresholds of the standard decoder (trained and tested for each coherence) are plotted against the thresholds of the decoders trained only at 100% coherence (and tested at every other coherence). The median threshold difference was not statistically significantly different to zero (p = 0.826, Wilcoxon Rank Sign test). B: Mean difference in accuracy between standard decoders and decoders trained only at 100% coherence, plotted at each coherence level, showing that performance was similar at all levels of coherence not just near threshold (1-way repeated measures ANOVA F_4_ = 1.61, p = 0.18). Error bars represent the standard error of the mean. No individual data point was significantly different from 0 (p > 0.05). C: Normalized decoder weights of the per coherence decoders plotted against the 100% coherence decoders. Each point represents the weight of an individual unit, filled circles represent weights that are statistically different at the 100% and near threshold conditions. The weights of the two decoder types were strongly correlated (Spearman’s ρ = 0.711, p < 0.001). D-E: Effects of correlations on decoders trained at 100% coherence. D: The changes in thresholds for the 100% coherence trained decoders and the per coherence trained decoders was very similar when removing correlations (Spearman’s ρ = 0.938, p < 0.001). As in Figure 4F, the change in threshold was calculated as the standard threshold minus the no correlations threshold. E: The changes in threshold for the two types of decoders was also very similar when ignoring correlations (Spearman’s ρ = 0.715, p = 0.001). Similar to Figure 4F, the change in threshold was calculated as the standard threshold minus the ignore correlations threshold.

To further examine if the decoders that were optimized at 100% coherence are equivalent to the decoders optimized on a per coherence basis, we tested if the effect of removing or ignoring correlations was the same for these two types of decoder. We reasoned that if these two types of decoder were using the same decoding strategy, they should show similar effects when removing or ignoring correlations. We first compared the effects of removing correlations at 100% coherence to the same effects performed at a threshold correlation (Figure 5D). First, as expected, the majority of the data points were in the top right quadrant, indicating that removing correlation had the same effect (a decrease in threshold) regardless of whether training was done with the 100% coherence trials, or at the coherence of which the decoder was tested. Second, these data points were highly correlated (Spearman’s ρ = 0.938, p < 0.001), indicating that the effects were invariant to the two training methods. We also compared the effects of ignoring correlations for decoders that were trained at 100% coherence with decoders that were trained on a per coherence basis (Figure 5E). We found that the change in threshold for the decoders that were trained at 100% coherence was highly correlated with per coherence trained decoders when correlations were ignored (Figure 5E, Spearman’s ρ = 0.715, p = 0.001). Altogether, our data suggest that the optimal linear decoding strategy is invariant to the level of motion coherence.

### Decoder weights are largely determined by left-right selectivity, not preferred direction

Finally, we examined how well an individual unit’s direction selectivity predicts its decoding weight. We considered two types of direction selectivity: left-right selectivity (Equation 2), and the vertical meridian offset, which we defined as difference between the preferred direction of the unit (e.g. Figure 1, left panels) and the vertical meridian. For example, a unit that prefers motion to the right would have a vertical meridian offset of +90°, whereas one that prefers leftwards motion would be −90°, and units that prefer upwards or downwards motion would be 0°. While left-right selectivity and vertical meridian offset are related_(Spearman’s ρ = 0.769, p < 0.001), they are not necessarily equivalent. For example, it is possible for a unit that prefers rightwards motion (e.g. 0°) to have weaker left-right selectivity than a unit that prefers motion at 45° due to differences in in firing rates and tuning bandwidths. Furthermore, even though left-right selectivity should be a strong predictor of a unit’s weight in a left-right decoding task, the correlation structure of real neuronal populations may result in units whose preferred axis of motion lies along the left-axis being more informative than those that are offset. Therefore, we tested if the decoder weights were determined simply by left-right selectivity, or if they were also influenced by the preferred direction of the units.

Figure 6A-C shows the normalized decoder weights at 100% coherence plotted against the left-right selectivity for 3 example populations, using both the standard decoders (correlations present) and the decoders trained on trial shuffled data (no correlations). Populations typically showed a clear relationship between left-right selectivity and weights for both types of decoder, but the shuffled decoder weights showed a much tighter relationship. Across all populations, unit weights were highly correlated with left-right selectivity (Figure 6D, left panel, Spearman’s ρ = 0.841, p < 0.001), especially for decoders trained on trial shuffled data (Figure 6D, right panel, Spearman’s ρ = 0.954, p< 0.001). This indicates that while left-right selectivity is a key factor affecting the decoding weight, the weight may be slightly offset to optimize decoding performance when correlations are present.

In contrast, the strength of the correlation between unit weight and vertical meridian offset was much weaker (Figure 6E, left panel, Spearman’s ρ = 0.594, p < 0.001), even for decoders trained on trial shuffled data (Figure 6E, right panel, Spearman’s ρ = 0.738, p < 0.001). Furthermore, there was no significant correlation between unit weight and vertical meridian offset when controlling for left-right selectivity (partial correlation, Spearman’s ρ = −0.159, p = 0.06). In the summary, these results show that decoder weights are largely determined by left-right selectivity rather than the preferred direction of the unit.

**Figure 6:**
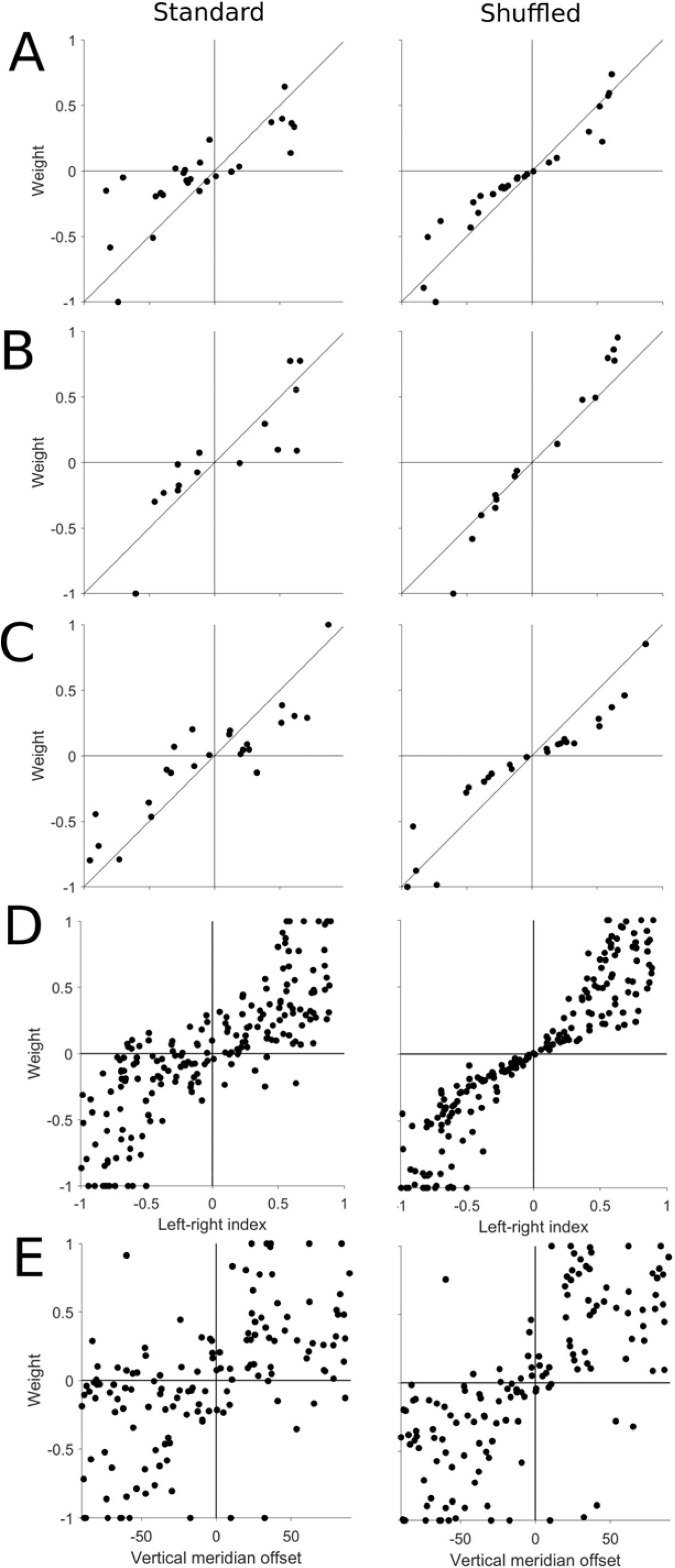
Decoders weights and direction selectivity. A-C Three example populations showing the normalized decoder weights plotted against the left-right selectivity for the standard decoders (left column, correlations present) and the decoders trained on shuffled data (right column, no correlations present). D: Decoder weights for all populations plotted against left-right selectivity, showing a strong relationship (left), especially when spiking correlations are removed (right). E: Decoder weights for all populations plotted against vertical meridian offset, showing a much weaker relationship (left), even when spiking correlations were not present (right).

## Discussion

In this paper, we have presented the most extensive recordings to date of population activity in areas MT and MST for random dot motion embedded in noise. We found that correlations impaired decoding performance, but decoders which learnt the correlation structure performed better than those that ignored correlations. We also found that decreases in motion coherence led to increases in correlations that were independent of changes in spike counts. Despite these changes in correlations with coherence, decoders that were trained using only 100% coherence stimuli performed as well as decoders that were optimized for each level of coherence. Finally, we showed the decoder weights were primarily dependent on the left-right selectivity of the unit, not the preferred direction. These results provide valuable new insights into the correlations and information contained in real populations of neurons for opposite directions of motion discrimination tasks.

### Stronger correlations in response to weaker motion signals

We found that the mean spike count correlation (rSC) in MT and MST in response to random dot stimuli ranged from 0.17 (100% coherence) to 0.26 (10% coherence). The strength of the r_SC_ correlations at 100% coherence was similar to previous studies that recorded pairs of nearby neurons in MT (0.12: Zohary et al., 1994; 0.2: Bair et al., 2001; 0.13: Cohen and Newsome, 2008; 0.1: Huang and Lisberger, 2009), even though these studies were conducted in awake animals, whereas ours were anesthetized. In the primary visual cortex, correlations are higher in anesthetized preparations compared to awake animals (Ecker et al., 2014). These differences seem to arise from changes in “network state”, which is correlated with the low frequency range of the local field potential (Ecker et al., 2014). We did not take any measure of network states, but given the similarity of our measurements of rSC with those in awake animals, it is possible that anesthesia does not change the magnitude of r_SC_ values in MT and MST.

Furthermore, our finding that r_SC_ was higher in response to weaker motion signals compared to stronger motion signals complemented the findings of Bair et al. (2001), who had reported the same result for pairs of neurons that prefer similar directions in awake animals. However, their results were based on a sample of 29 pairs, therefore our analysis substantially builds on this by examining over 1000 pairs of units with a range of differences in direction selectivity (Figure 2B,C). Furthermore, we have shown that these results were not a consequence of changes in spike counts (Cohen and Kohn, 2011). While it is known that anesthesia can affect correlation structure in primary visual cortex by increasing r_SC_ values for high firing rate pairs and pairs with similar tuning (Ecker et al., 2014), our observation that low coherence stimuli produce higher correlations was not dependent on firing rate (e.g. Figure 3B) or signal correlation (Figure 3E,F), and additionally. The finding that weaker stimulus strength produces greater rSC values is also in agreement with the effects of changes in contrast of sinusoidal gratings on r_SC_ measurements in the primary visual cortex of macaques (Smith and Kohn, 2008), where lower contrast gratings (which elicit lower spikes rates) produced higher correlations. These results, in combination with ours, suggest that weaker stimulus strengths may generally result in higher neuronal correlations in the visual cortex.

Since the noise in our stimulus was generated randomly for each stimulus presentation, it is possible that the neuronal correlations arise because of the variability in the stimulus, rather than having a neural origin. However, Bair et al. (2001) found no significant difference in correlations for randomly generated stimuli and identical presentations of the same stimuli. Furthermore, Britten et al. (1993) compared the spiking activity of single neurons in response to both randomly and statically generated stimulus noise found no difference in spiking variability, suggesting that the variability of responses (and therefore correlations) is not related to the random nature of the stimulus. Nonetheless, whether or not the correlations were, in part, driven by the stimulus, does not impact the analyses and conclusions of this study.

### Effects of correlations in population decoding

Our finding that correlations impair neural decoding is in agreement with previous works (e.g. Zohary et al., 1994), with the simplest explanation being that noise cannot be averaged out in the presence of correlations. Our results are also compatible with studies of attention, which show that attention decreases neuronal correlations and improves stimulus feature decoding (Cohen and Maunsell, 2009, 2011; Mitchell et al., 2009). On the other hand, recent work has also demonstrated that, both theoretically and empirically, the presence of trial-to-trial correlations can improve population decoding (Abbott and Dayan, 1999; Sompolinsky et al., 2001; Averbeck et al., 2006; Shamir and Sompolinsky, 2006; Ecker et al., 2011; Graf et al., 2011; Kohn et al., 2016; Zylberberg et al., 2016; Zavitz et al., 2017). Whether the correlation structure helps or hinders population decoding may be dependent on the type of task performed by the decoder (Ecker et al., 2011). Studies that have found that population decoding performance was improved by the presence of correlations usually readout a continuous estimate of a stimulus parameter (Graf et al., 2011; Zavitz et al., 2016; Zylberberg et al., 2016), which is comparable with a fine discrimination task in which subjects make judgements between small differences in stimulus attributes. However, in our study, the decoder made a binary choice between two opposites directions in noisy conditions, suggesting the correlation structure of neurons in MT/MST may be not be beneficial for population coding in a 2AFC opposite direction task. The correlation structure of populations of sensory neurons will depend, at least to some degree, on the task that the animal is performing (Cohen and Newsome, 2008), so it is possible that the correlations may be helpful, or less harmful to performance, when the animal is engaged in the task.

It should be noted that the effects of correlations can depend on population size (Lin et al., 2015; Kohn et al., 2016), in which small populations (<100) that undersample the full population can lead to the false conclusion that presence of correlations improves decoding, whereas larger samples of the full population demonstrate that correlations impair decoding. However, we found that the presence of correlations *impair* decoding in *small* populations, and to our knowledge there is no study that demonstrates that this effect would be reversed for larger populations. The question of how decoding performance scaled with the number of neurons in the population is an important question. We found that population size had no significant effect on population threshold, but because of the heterogeneity of our populations, our data can only provide limited insights into this issue. Addressing this question in full will require larger populations that cover the full range of preferred directions and selectivities.

We also found ignoring the correlation structure had a small but significant impact on decoding performance, implying that downstream neurons have to know the correlations between neurons in order to extract all available information. However, knowing or learning correlations is not unrealistic, as correlations were relatively consistent between coherence, implying that only one set of correlations has to be known for near-optimal decoding. Indeed, applying a linear decoder trained with the 100% coherence stimuli was no less effective than decoders trained per coherence (Figure 5A). The fact that neurons that carried the most information at 100% coherence will likely be the most informative at lower coherences (Chaplin et al., 2017) also meant that optimal decoding weights can be predicted fairly accurately at lower coherences from those trained at 100% (Figure 5B). This also meant that the effects observed when weights were optimized per coherence, i.e. the effects of removing and ignoring correlations, were also present when the 100% coherence weights were used universally (Figure 5D-C). Altogether, these results contributed to the reasons why weights trained at 100% coherence can be applied to lower coherences with little loss in performance.

Interestingly, we observed that the populations that were most impaired by the presence of correlations showed the least improvement from learning the correlations, and populations which showed the most improvement from when correlations were considered were least affected correlations were removed (Figure 4D). This result may appear to be counter-intuitive, however it has been shown computationally that both scenarios are possible (Averbeck et al., 2006).

### Optimal weighting of responses for decoding in opposite directions of motion task

Our results show that the optimal weights are dependent on the left-right selectivity of units rather than their preferred direction, with some optimizations to account for correlations. This reflects previous studies that suggest that perceptual learning can be best accounted for by optimizing weights via changes in feedforward connectivity (Law and Gold, 2009; Bejjanki et al., 2011), as the improvements sensory representation of stimulus features appear to be minimal (Schoups et al., 2001; Yang and Maunsell, 2004; Raiguel et al., 2006), particularly for area MT in a 2AFC opposite directions of motion task (Law and Gold, 2008). Essentially, such a learning process enables the neurons which carry the most task-relevant information to contribute the most to the decision, which is a similar process to training the decoder to optimize weights for left-right decoding in the present study. Therefore perceptual learning could be mediated by a process that learns a slightly different set of weights that are specific to the task, which in this case would be the left-right selectivities and correlation structure, not the preferred direction. Since the present study also demonstrates that decoders can be trained to perform the task at 100% coherence, and then apply same decoding strategy at lower coherences, and still perform relatively well, then perceptual learning could be mediated a simpler process than if the weights have to be refined substantially with respect to changes in coherences. Furthermore, this means that downstream areas that readout MT/MST neurons do not, in principle, need to first know the level of motion coherence, as the optimal linear readout is the same for all motion coherences. This also has important implications for modelling of neuronal populations for sensory readout in 2AFC tasks (Shadlen et al., 1996; Cohen and Newsome, 2009; Wimmer et al., 2015).

## References

Abbott LF, Dayan P (1999) The effect of correlated variability on the accuracy of a population code. Neural Comput 11:91–101.

Adibi M, McDonald JS, Clifford CWG, Arabzadeh E (2014) Population decoding in rat barrel cortex: optimizing the linear readout of correlated population responses. PLoS Comput Biol 10:e1003415.

Albright TD (1984) Direction and orientation selectivity of neurons in visual area MT of the macaque. J Neurophysiol 52:1106–1130.

Averbeck BB, Crowe DA, Chafee M V., Georgopoulos AP (2003) Neural activity in prefrontal cortex during copying geometrical shapes. Exp Brain Res 150:142–153.

Averbeck BB, Latham PE, Pouget A (2006) Neural correlations, population coding and computation. Nat Rev Neurosci 7:358–366.

Bair W, Zohary E, Newsome WT (2001) Correlated firing in macaque visual area MT: time scales and relationship to behavior. J Neurosci 21:1676–1697.

Bejjanki VR, Beck JM, Lu ZL, Pouget A (2011) Perceptual learning as improved probabilistic inference in early sensory areas. Nat Neurosci 14:642–650.

Berens P, Ecker AS, Gerwinn S, Tolias AS, Bethge M (2011) Reassessing optimal neural population codes with neurometric functions. Proc Natl Acad Sci 108:4423–4428.

Born RT, Bradley DC (2005) Structure and function of visual area MT. Annu Rev Neurosci 28:157–189.

Bourne JA, Rosa MGP (2003) Preparation for the in vivo recording of neuronal responses in the visual cortex of anaesthetised marmosets (Callithrix jacchus). Brain Res Protoc 11:168–177.

Brainard DH (1997) The psychophysics toolbox. Spat Vis 10:433–6.

Britten KH, Newsome WT, Shadlen MN, Celebrini S, Movshon JA (1996) A relationship between behavioral choice and the visual responses of neurons in macaque MT. Vis Neurosci13:87–100.

Britten KH, Shadlen MN, Newsome WT, Movshon JA (1992) The analysis of visual motion: a comparison of neuronal and psychophysical performance. J Neurosci 12:4745–4765.

Britten KH, Shadlen MN, Newsome WT, Movshon JA (1993) Responses of neurons in macaque MT to stochastic motion signals. Vis Neurosci 10:1157–1169.

Celebrini S, Newsome WT (1994) Neuronal and psychophysical sensitivity to motion signals in extrastriate area MST of the macaque monkey. J Neurosci 14:4109–4124.

Chaplin TA, Allitt BJ, Hagan MA, Price NSC, Rajan R, Rosa MGP, Lui LL (2017) Sensitivity of neurons in the middle temporal area of marmoset monkeys to random dot motion. J Neurophysiol 118:1567–1580.

Cohen MR, Kohn A (2011) Measuring and interpreting neuronal correlations. Nat Neurosci 14:811–819.

Cohen MR, Maunsell JHR (2009) Attention improves performance primarily by reducing interneuronal correlations. Nat Neurosci 12:1594–1600.

Cohen MR, Maunsell JHR (2011) Using neuronal populations to study the mechanisms underlying spatial and feature attention. Neuron 70:1192–1204.

Cohen MR, Newsome WT (2008) Context-dependent changes in functional circuitry in visual area MT. Neuron 60:162–173.

Cohen MR, Newsome WT (2009) Estimates of the contribution of single neurons to perception depend on timescale and noise correlation. J Neurosci 29:6635–6648.

de la Rocha J, Doiron B, Shea-Brown E, Josić K, Reyes A (2007) Correlation between neural spike trains increases with firing rate. Nature 448:802–6.

Duffy CJ, Wurtz RH (1991) Sensitivity of MST neurons to optic flow stimuli. I. A continuum of response selectivity to large-field stimuli. J Neurophysiol 65:1329–1345.

Ecker AS, Berens P, Cotton RJ, Subramaniyan M, Denfield GH, Cadwell CR, Smirnakis SM, Bethge M, Tolias AS (2014) State dependence of noise correlations in macaque primary visual cortex. Neuron 82:235–248.

Ecker AS, Berens P, Tolias AS, Bethge M (2011) The effect of noise correlations in populations of diversely tuned neurons. J Neurosci 31:14272–14283.

Gallyas F (1979) Silver staining of myelin by means of physical development. Neurol Res 1:203–209.

Goris RLT, Movshon JA, Simoncelli EP (2014) Partitioning neuronal variability. Nat Neurosci 17:858–865.

Graf AB, Kohn A, Jazayeri M, Movshon JA (2011) Decoding the activity of neuronal populations in macaque primary visual cortex. Nat Neurosci 14:239–245.

Huang X, Lisberger SG (2009) Noise correlations in cortical area MT and their potential impact on trial-by-trial variation in the direction and speed of smooth-pursuit eye movements. J Neurophysiol 101:3012–30.

Kanitscheider I, Coen-Cagli R, Pouget A (2015) Origin of information-limiting noise correlations. Proc Natl Acad Sci 112:E6973–E6982.

Kohn A, Coen-Cagli R, Kanitscheider I, Pouget A (2016) Correlations and neuronal population information. Annu Rev Neurosci 39:237–256.

Law C-T, Gold JI (2008) Neural correlates of perceptual learning in a sensory-motor, but not a sensory, cortical area. Nat Neurosci 11:505–513.

Law C-T, Gold JI (2009) Reinforcement learning can account for associative and perceptual learning on a visual-decision task. Nat Neurosci 12:655–663.

Leavitt ML, Pieper F, Sachs AJ, Martinez-Trujillo JC (2017) Correlated variability modifies working memory fidelity in primate prefrontal neuronal ensembles. Proc Natl Acad Sci 114:E2494–E2503.

Lin I-C, Okun M, Carandini M, Harris KD (2015) The nature of shared cortical variability. Neuron 87:644–656.

Lui LL, Rosa MGP (2015) Structure and function of the middle temporal visual area (MT) in the marmoset: Comparisons with the macaque monkey. Neurosci Res 93:62–71.

Maunsell JHR, Van Essen DC (1983) Functional properties of neurons in middle temporal visual area of the macaque monkey. I. Selectivity for stimulus direction, speed, and orientation. J Neurophysiol 49:1127–1147.

Mitchell JF, Sundberg KA, Reynolds JH (2009) Spatial attention decorrelates intrinsic activity fluctuations in macaque area V4. Neuron 63:879–88.

Moreno-Bote R, Beck J, Kanitscheider I, Pitkow X, Latham P, Pouget A (2014) Information-limiting correlations. Nat Neurosci 17:1410–1417.

Newsome WT, Britten KH, Movshon JA (1989) Neuronal correlates of a perceptual decision. Nature 341:52–54.

Palmer SM, Rosa MGP (2006) A distinct anatomical network of cortical areas for analysis of motion in far peripheral vision. Eur J Neurosci 24:2389–2405.

Paxinos G, Watson C, Petrides M, Rosa MGP, Tokuno H (2012) The marmoset brain in stereotaxic coordinates. Academic Press.

Pesaran B, Pezaris JS, Sahani M, Mitra PP, Andersen RA (2002) Temporal structure in neuronal activity during working memory in macaque parietal cortex. Nat Neurosci 5:805–811.

Raiguel S, Vogels R, Mysore SG, Orban GA (2006) Learning to see the difference specifically alters the most informative V4 neurons. J Neurosci 26:6589–6602.

Ringach DL, Shapley RM, Hawken MJ (2002) Orientation selectivity in macaque V1: diversity and laminar dependence. J Neurosci 22:5639–51.

Rosa MGP, Elston GN (1998) Visuotopic organisation and neuronal response selectivity for direction of motion in visual areas of the caudal temporal lobe of the marmoset monkey (Callithrix jacchus): middle temporal area, middle temporal crescent, and surrounding cortex. J Comp Neurol 393:505–527.

Ruff DA, Cohen MR (2016) Attention increases spike count correlations between visual cortical areas. J Neurosci 36:7523–7534.

Saito HA, Yukie M, Tanaka K, Hikosaka K, Fukada Y, Iwai E (1986) Integration of direction signals of image motion in the superior temporal sulcus of the macaque monkey. J Neurosci 6:145–57.

Schoups A, Vogels R, Qian N, Orban GA (2001) Practising orientation identification improves orientation coding in V1 neurons. Nature 412:549–553.

Shadlen MN, Britten KH, Newsome WT, Movshon JA (1996) A computational analysis of the relationship between neuronal and behavioral responses to visual motion. J Neurosci 16:1486–1510.

Shamir M (2014) Emerging principles of population coding: In search for the neural code. Curr Opin Neurobiol 25:140–148.

Shamir M, Sompolinsky H (2006) Implications of neuronal diversity on population coding. Neural Comput 18:1951–86.

Smith M a, Kohn A (2008) Spatial and temporal scales of neuronal correlation in primary visual cortex. J Neurosci 28:12591–603.

Solomon SS, Chen SC, Morley JW, Solomon SG (2015) Local and global correlations between neurons in the middle temporal area of primate visual cortex. Cereb Cortex 25:3182–3196.

Solomon SS, Tailby C, Gharaei S, Camp AJA, Bourne JA, Solomon SG (2011) Visual motion integration by neurons in the middle temporal area of a New World monkey, the marmoset. J Physiol 23:5741–5758.

Sompolinsky H, Yoon H, Kang K, Shamir M (2001) Population coding in neuronal systems with correlated noise. Phys Rev E 64:051904.

Tanaka K, Hikosaka K, Saito H a, Yukie M, Fukada Y, Iwai E (1986) Analysis of local and wide-field movements in the superior temporal visual areas of the macaque monkey. J Neurosci 6:134–44.

Tolhurst DJ, Movshon JA, Dean AF (1983) The statistical reliability of signals in single neurons in cat and monkey visual cortex. Vision Res 23:775–85.

Wimmer K, Compte A, Roxin A, Peixoto D, Renart A, de la Rocha J (2015) Sensory integration dynamics in a hierarchical network explains choice probabilities in cortical area MT. Nat Commun 6:6177.

Yang T, Maunsell JHR (2004) The effect of perceptual learning on neuronal responses in monkey visual area V4. J Neurosci 24:1617–26.

Yu H-H, Rosa MGP (2010) A simple method for creating wide-field visual stimulus for electrophysiologyH: Mapping and analyzing receptive fields using a hemispheric display. J Vis 10:1–16.

Zavitz E, Yu H-H, Rosa MGP, Price NSC (2017) Correlated variability in the neurons with the strongest tuning improves direction coding. Cereb Cortex 1–12.

Zavitz E, Yu H-H, Row EG, Rosa MGP, Price NSC (2016) Rapid adaptation induces persistent biases in population codes. J Neurosci 36:4579–4590.

Zeki SM (1974) Functional organization of a visual area in the posterior bank of the superior temporal sulcus of the rhesus monkey. J Physiol 236:549–573.

Zohary E, Shadlen MN, Newsome WT (1994) Correlated neuronal discharge rate and its implications for psychophysical performance. Nature.

Zylberberg J (2017) Untuned but not irrelevant: A role for untuned neurons in sensory information coding. bioRxiv 1–18.

Zylberberg J, Cafaro J, Turner MH, Shea-Brown E, Rieke F (2016) Direction-selective circuits shape noise to ensure a precise population code. Neuron 89:369–383.

